# Lipidoid nanoparticles increase ATP uptake into hypoxic brain endothelial cells

**DOI:** 10.1101/2022.04.07.487513

**Authors:** Purva Khare, James F. Conway, Devika S Manickam

## Abstract

Lipidoid nanoparticles (LNPs) are clinically successful carriers for nucleic acid delivery to liver and muscle targets. Their ability to load and deliver small molecule drugs has not been reported yet. We propose that the delivery of adenosine triphosphate (ATP) to brain endothelial cells (BECs) lining the blood-brain barrier may increase cellular energetics of the injured BECs. We formulated and studied the physicochemical characteristics of ATP-loaded LNPs using the C12-200 ionizable cationic lipid and other helper lipids. Polyethylene glycol-dimyristoyl glycerol (PEG-DMG), one of the helper lipids, played a crucial role in maintaining colloidal stability of LNPs over time whereas the inclusion of both ATP and PEG-DMG maintained the colloidal stability of LNPs in the presence of serum proteins. ATP-LNPs formulated with PEG-DMG resulted in a 7.7- and 6.6-fold increased uptake of ATP into normoxic and hypoxic BECs, respectively. Altogether, our results demonstrate the potential of LNPs as a novel carrier for the delivery of small molecular mass actives to BECs—a CNS target.

**Highlights:**

- LNPs were formulated with ATP, a small molecule drug
- PEG-DMG plays a critical role in maintaining particle stability over tim
- ATP and PEG-DMG play a critical role in maintaining particle stability in 10% serum
- ATP-LNPs were internalized by normoxic and hypoxic brain endothelial cells (BECs)
- LNP delivery to BECs broadens its applicability to CNS targets

**Graphical Abstract:** 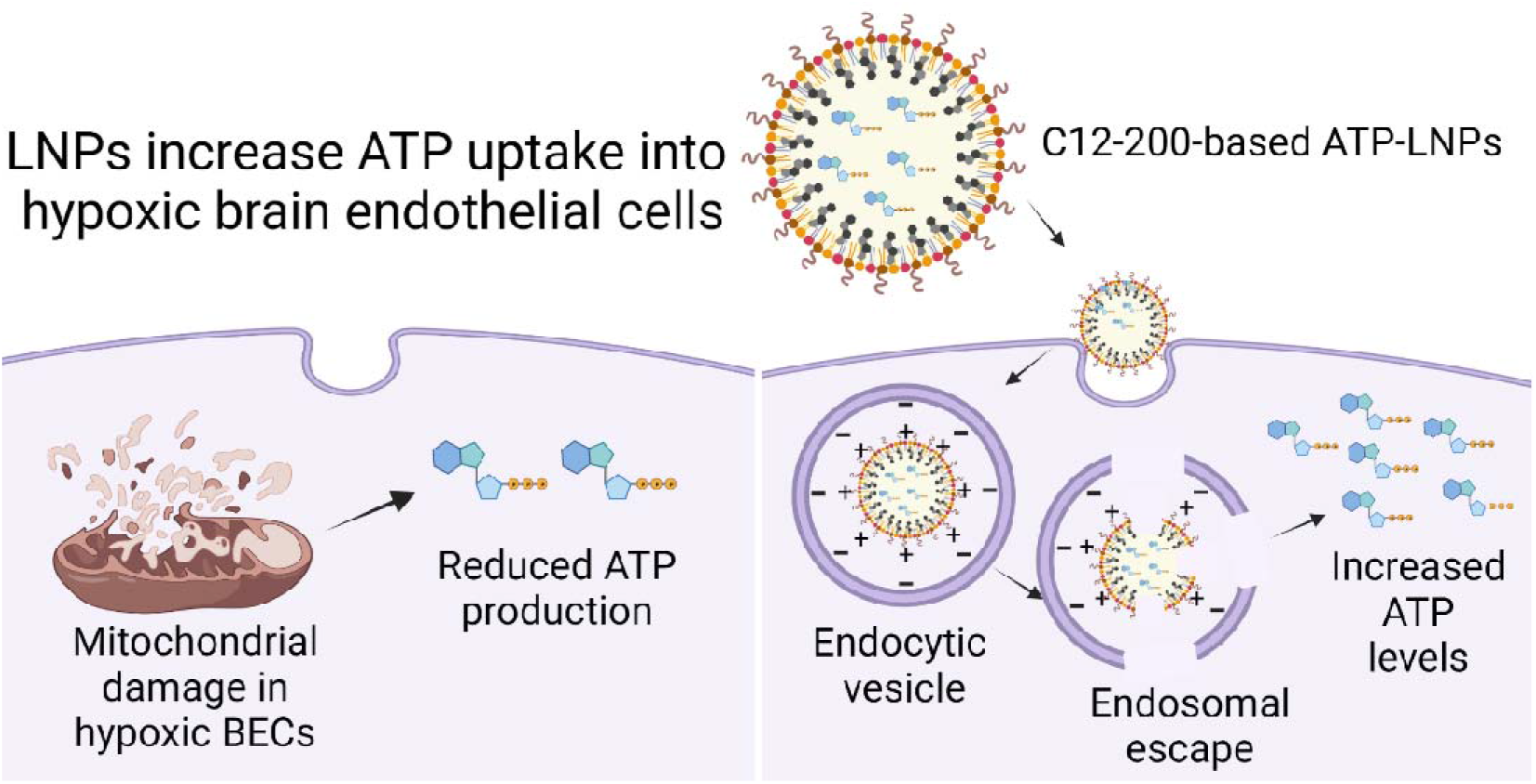

## Introduction

Lipidoid nanoparticles (LNPs) have revolutionized the field of non-viral delivery systems (1, 2) and are mRNA carriers in the Pfizer-BioNTech (Comirnaty) and Moderna (Spikevax) COVID-19 mRNA vaccines (3, 4). Onpattro was the first FDA-approved siRNA-LNP drug for the treatment of polyneuropathies caused by hereditary transthyretin amyloidosis (5). LNPs have been extensively studied for the delivery of large molecules like polypeptides and nucleic acids (6-11), however, their delivery potential for small molecule drugs remains unexplored. We propose that delivery of adenosine triphosphate (ATP) to the brain endothelial cells (BECs) lining the blood-brain barrier (BBB) is an effective strategy to increase the cell viability of BECs post-cerebral ischemia/reperfusion injury (stroke). Rescuing ATP levels in the BECs lining the BBB can lead to protection of its metabolic function and structural integrity, thereby limiting long-term damage to the brain tissue (12-15). Delivery of ATP, an anionic molecule is challenging due to its short half-life (∼ 10 minutes) and its limited permeability across cellular membranes (16, 17).

Liposomes have been extensively studied for loading ATP and have demonstrated increased energetics in ischemic liver, cardiac and brain tissues (18-25). While liposomes are an established delivery platform, the thin film hydration method results only in a modest loading of 5 mol% ATP (18). Further process modifications led to the development of reverse phase evaporation and freezing-thawing methods resulting in a maximum entrapment of 36 to 38 mol% ATP (18). LNPs have reported high siRNA and mRNA encapsulation efficiencies in the order of ca. 80-98% (26). Based on their higher encapsulation efficiencies and successful clinical performance, we sought to determine if LNPs can be formulated for the delivery of ATP.

LNPs are composed of an ionizable cationic lipid and other helper lipids like cholesterol, polyethylene glycol-dimyristoyl glycerol (PEG-DMG) and distearylphosphatidylcholine (DSPC) (27, 28). LNPs possess numerous advantages such as high encapsulation efficiency of the cargo, protection of the cargo from enzymatic degradation, improved pharmacokinetics, and lower toxicity and immunogenicity (5, 29-34). Ionizable cationic lipids allow superior cargo encapsulation of anionic cargo along with their efficient intracellular release due to endosomal escape (35). Seminal studies have demonstrated the superior potency of C12-200 in delivering siRNA to liver targets in rodents and non-human primates (36-38).

We have previously demonstrated that C12-200-based LNPs increased siRNA uptake into primary neurons, a tenacious target cell (39). In this work, we again used C12-200 to formulate ATP-LNPs. We studied the physicochemical characteristics and compared the colloidal and serum stability of blank- vs. ATP-loaded LNPs using dynamic light scattering. We then compared the morphology of ATP-LNPs and the widely studied siRNA-LNPs using transmission electron microscopy. Following that, we studied the cellular uptake of ATP-LNPs in two different models of hypoxia-subjected BECs: a human brain microvascular endothelial cell line (hCMEC/D3) and primary human brain microvascular endothelial cells (HBMECs) using fluorescence microscopy and flow cytometry.

## Experimental section

### Materials

Cholesterol (8667) was obtained from Sigma-Aldrich (St. Louis, MO). 1,2-Dimyristoyl-rac– glycero-3-methoxypolyethylene glycol-2000 (PEG-DMG) (8801518) and 1,2-distearoyl-sn-glycero-3-phosphocholine (DSPC) (850365P) were procured from Avanti Polar Lipids (Alabaster, AL). The ionizable cationic lipid, 1,1‘-((2-(4-(2-((2-(bis (2 hydroxy dodecyl) amino) ethyl) (2hydroxydodecyl) amino) ethyl) piperazin1yl) ethyl) azanediyl) bis(dodecan-2-ol) (C12-200) was a generous gift from Dr. Muthiah Manoharan, Alnylam Pharmaceuticals (Cambridge, MA). Lipofectamine RNAiMAX (13778075) and Alexa Fluor 647-labeled ATP (AF647-ATP) (A22362) were procured from Thermo Fisher Scientific (Waltham, MA). Phosphate-buffered saline (PBS) and heat-inactivated fetal bovine serum (FBS) were obtained from Hyclone Laboratories (Logan, UT). Primary Human Brain Microvascular Endothelial Cells (HBMECs) (ACBRI 376) were purchased from Cell Systems (Kirkland, WA) at passage number 3 (P3). HBMECs at P3 and P12 were used in all experiments. The complete cell culture kit (The System; CSS-A101) for growing and maintaining HBMECs was purchased from Cell Systems (Kirkland, WA). The System consisted of a medium component, growth supplement component, passage reagent group (PRG) component, cell freezing medium and an attachment factor. Human cerebral microvascular endothelial cell line (hCMEC/D3) (102114.3C) at passage number (P) 25 was obtained from Cedarlane Laboratories (Burlington, Ontario). hCMEC/D3 cells maintained between P25 and P35 were used in all experiments. All reagents were used as received unless stated otherwise. Hoechst 33342 (H3570) was procured from Fisher Scientific (Waltham, MA).

### Preparation of ATP-loaded LNPs (ATP-LNPs)

LNPs were formulated according to previously reported methods (40). Briefly, DSPC, cholesterol, PEG-DMG and the ionizable cationic lipidoid C12-200 were dissolved at a molar ratio of 50/10/38.5/1.5 in ethanol. The representative formulation scheme for the preparation of ATP- and siRNA-loaded LNPs are shown in ***Table 1*** and Supplementary table 1, respectively. For ATP-LNPs, a 33.73 mg/mL solution of ATP was prepared in 1*x* PBS pH 7.4. We have previously studied the effects of mixing speeds on drug loading by comparing ‘slow mixing’ and ‘fast mixing’ protocols based on the speed used during the addition of the lipid/ethanolic phase to the aqueous phase (39). The ‘fast mixing’ protocol resulted in increased loading of siRNA into the LNPs and so we used the same for preparing ATP-LNPs. For the ‘fast mixing’ protocol, the ethanolic phase was added in a dropwise manner to the aqueous phase containing ATP under continuous vortexing for 30 seconds. This was achieved by maintaining the Fisher benchtop vortexer knob at position ‘7’. ATP-LNP formulations were prepared at a final ATP concentration of 50 mM. For fluorescence microscopy and flow cytometry analysis, ATP-LNPs were volumetrically spiked 1:100 or 1:1000 with AF647-ATP.

**Table 1.** Representative formulation scheme for ATP-LNPs.

| Component | w/w% in final LNP mixture | Stock concentration (mg/mL) | Volume of the stock solution required (μL) |
| --- | --- | --- | --- |
| **Ethanolic phase** |  |  |  |
| C12-200 | 50 | 1 | 12.5 |
| Cholesterol | 38.5 | 1 | 3.1 |
| PEG-DMG | 1.5 | 0.5 | 13.6 |
| DSPC | 10 | 0.5 | 16.9 |
| Ethanol | - | - | 78.9 |
| **Total ethanolic phase volume** |  |  | 125 |
| **Aqueous phase** |  |  |  |
| ATP in 1x PBS | 50 mM | 33.73 | 375 |
| **Total aqueous phase volume** |  |  | 375 |
| **Final volume of LNPs prepared (μL)** |  |  | 500 |

### Dynamic Light Scattering

We studied the colloidal stability of LNPs by measuring their average particle diameters, dispersity indices and zeta potentials (particle surface charge) of the blank and ATP-loaded LNPs prepared with (+) and without (-) PEG-DMG using dynamic light scattering (DLS) on a Malvern Zetasizer Pro (Malvern Panalytical Inc., Westborough, PA). We measured particle diameters and surface charge over a period of seven days with intermittent storage at 2-8 ºC. We also studied the colloidal stability of LNPs in 1*x* PBS and in 10% fetal bovine serum upon storage at 37 ºC for 24 h to determine the effect of serum proteins/cell culture conditions. LNPs were diluted to a final ATP concentration of 5 mM in 1*x* PBS pH 7.4 and in deionized water for particle diameter and zeta potential measurements, respectively. Data are represented as average ± standard deviation (SD) of triplicate measurements. The data are representative of at least five-six independent experiments (measurement error <5%).

### Transmission Electron Microscopy

LNPs were prepared for negative stain electron microscopy by applying a 3 µL volume to a freshly glow-discharged continuous carbon on a copper grid and staining with a 1% uranyl acetate solution. Grids were inserted into a Thermofisher TF20 electron microscope (Thermofisher Scientific, Waltham, Massachusetts, USA) equipped with a field emission gun and imaged on a TVIPS XF416 CMOS camera (TVIPS GmbH, Gauting, Germany) to visualize nanoparticle dimensions. The microscope was operated at 200 kV and contrast was enhanced with a 40 µm objective aperture. Electron micrographs were collected at a nominal 19,000x magnification with a post-column magnification of 1.3x corresponding to a calibrated pixel size of 5.6 Ångstroms at the sample. Images were acquired with TVIPS *Emplified* software using movie mode for drift correction.

### Cell culture

hCMEC/D3 brain endothelial cell line was maintained in tissue culture flasks pre-coated with 0.15 mg/mL rat collagen I in a humidified incubator at 37 ºC and 5% CO_2_ (Isotemp, Thermo Fisher Scientific). hCMEC/D3 cells were maintained in complete growth medium comprising of endothelial cell basal medium (EBM-2) supplemented with fetal bovine serum/FBS (5%), ascorbic acid (5 µg/mL), hydrocortisone (1.4 µM), 10 mM HEPES (pH 7.4), bFGF (1 ng/mL), chemically defined lipid concentrate (0.01%) and penicillin (100 units/mL)-streptomycin (100 µg/mL) mixture. The complete growth medium was changed every other day until the cells formed a confluent monolayer. Cells were passaged after the formation of a confluent monolayer with 1*x* TrypLE express (Gibco, Denmark) after washing with 1*x* PBS. Primary HBMECs were maintained in tissue culture flasks pre-treated with the cell attachment factor. HBMECs were supplemented with the classic culture medium containing 1% v/v cell culture boost (4Z0-500). The complete growth medium was replenished every 48 h. Cells were passaged after the formation of a confluent monolayer with different passage group reagents (PRG) (4Z0-800). Cells were washed with PRG 1 (EDTA-dPBS solution), detached with PRG 2 (Trypsin/ EDTA-dPBS solution) and suspended in PRG 3 (Trypsin Inhibitor-dPBS solution). The cell suspension was centrifuged at 900*x*g for 10 min at 4 ºC. The supernatant was decanted, and the cell pellet was suspended in fresh classic culture medium for plating. For studies using cells maintained under normoxic conditions, they were supplemented with the complete growth medium and maintained in a humidified incubator at 37 ºC and 5% CO_2_. For studies under hypoxic conditions, cells were supplemented with the oxygen-glucose deprived (OGD) medium (14) and were maintained in a Billups-Rothenberg chamber pre-flushed with 5% carbon dioxide, 5% hydrogen and 90% nitrogen at maintained at 37 ± 0.5 °C.

### Uptake of LNPs into hCMEC/D3 cells using flow cytometry

hCMEC/D3 cells were seeded in a clear, 0.15 mg/mL rat collagen I and Poly-D-Lysine coated flat bottom 24-well plate (Azer Scientific, Morgantown, PA) at a density of 150,000 cells/well in complete growth medium until confluency in a humidified incubator at 37 ºC and 5% CO_2_. ATP-LNPs were prepared as mentioned in **Table 1** by additionally spiking ATP with AF647-ATP at a 100:1 ratio. AF647-ATP spiked ATP-LNPs were diluted using complete growth medium (normoxic) and OGD medium (hypoxic) to result in a final ATP concentration of 15 mM/well. The treatment mixture contained either free ATP or ATP-LNPs (+/- PEG-DMG) or Lipofectamine-ATP complexes. For cells exposed to hypoxic conditions, we pre-incubated them in OGD medium in the Billups-Rothenberg chamber for 24 h before treatment with LNPs or free ATP. The cells were incubated with 350 µL/well of the treatment mixture for 24 h after which the cells were washed with 200 µL of 1*x* PBS, detached with 100 µL Trypsin-EDTA solution and collected in microcentrifuge tubes. The cell suspension was centrifuged at 900*x*g for 10 minutes and the resulting cell pellet was resuspended in 1 mL of 2% FBS in PBS solution for flow cytometry analysis. Samples were analyzed on a Attune NxT Acoustic Focusing Cytometer (Singapore) equipped with Attune NxT software. AF647 fluorescence was detected at excitation and emission wavelengths of 650 nm and 665 nm, respectively. Dot plots were obtained from the Attune NxT software as percentages of AF647 positive cells AF647 (+) after gating out the autofluorescence in the unstained controls. Data are presented as -fold increase in the ATP uptake normalized to the untreated cells. Data are presented as mean ± SD of n=4 samples.

### Uptake of LNPs into HBMECs using fluorescence microscopy

HBMECs were seeded in a cell attachment factor-treated clear, flat bottom Poly-D-Lysine coated 24-well plate (Azer Scientific, Morgantown, PA) at a density of 150,000 cells/well in complete growth medium and were allowed to attain confluency in a humidified incubator at 37 °C and 5% CO_2_. ATP-LNPs were prepared by spiking ATP using AF647-ATP at a 1000:1 ratio. AF647 ATP-LNPs were diluted in the complete growth medium to result in a final ATP concentration of 5 mM/well. The treatment groups consisted of either free ATP or ATP-LNPs (+/- PEG-DMG) or Lipofectamine-ATP complexes. Cells were incubated with 350 µL/well of the treatment mixture for 24 h after which the treatment mixture was replaced with 1 mL phenol red free complete growth medium after washing the cells with 200 µL of 1*x* PBS. Cell nuclei were stained by incubating the cells with 10 µg/mL Hoechst dye for 10 minutes followed by washing with 200 µL of 1*x* PBS. Untreated cells were used as the negative control. Olympus IX 73 epifluorescent inverted microscope (Olympus, Center Valley, PA) was used to image the cells and AF647 signals were detected at excitation and emission wavelengths of 650 nm and 665 nm, respectively whereas Hoechst signal were detected at excitation and emission wavelengths of 350 nm and 461 nm, respectively. AF647 signals were normalized to untreated cells using the CellSens Software.

### Statistical analysis

Values are expressed as mean ± standard deviation (SD), wherever noted. Statistical analysis was performed using either paired t-test or one-way or two-way ANOVA using GraphPad Prism 9 (GraphPad Software, San Diego, CA). Comparative statistical analysis was performed using Tukey’s and Šídák’s multiple comparisons tests using two-way ANOVA. Bonferroni’s and Tukey’s multiple comparisons test was used for comparative analysis using one-way ANOVA. Alpha was set to 0.05.

## Results

### Physicochemical characterization and colloidal stability of ATP-LNPs

Physicochemical characteristics of nanoparticles govern their colloidal stability and *in vivo* biological efficacy (7, 41). Particle diameter and surface charge (zeta potential) are crucial determinants of colloidal stability. Cellular uptake of ionizable cationic lipid-based LNPs is largely dependent on their particle diameter homogeneity (42, 43). Colloidal particles like LNPs are known to aggregate resulting in larger particle diameters and polydisperse samples over time (44). We have previously demonstrated that siRNA loading further stabilized LNP particle diameters against aggregation (39). PEG-DMG acts as a steric stabilizer and is known to stabilize nanoparticles by hindering particle-particle aggregation resulting in monodisperse, smaller particle diameters (45, 46). Therefore, we compared the colloidal stability of blank and ATP-LNPs prepared in the presence or absence of PEG-DMG (+/- PEG-DMG) over a period of seven days upon interim storage at 2-8°C. We also measured LNP surface charge by measuring their zeta potential following the same storage regime.

Blank LNPs (+PEG-DMG) showed a particle diameter of ∼78 nm that increased to ∼167 nm whereas blank LNPs (-PEG-DMG) had a particle diameter of ∼895 nm that increased to ∼2 μm over a period of seven days **(Figure 1a)**. ATP-LNPs (+PEG-DMG) had an initial particle diameter of ∼83 nm that increased to ∼162 nm whereas ATP-LNPs (-PEG-DMG) showed a particle diameter of ∼995 nm post-preparation that increased to ∼3 μm after seven days **(Figure 1a)**. The increase in particle diameters among each group over a period of seven days was found to be non-significant using two-way ANOVA. However, the particle diameters of blank and ATP-LNPs prepared without PEG-DMG (-PEG-DMG) were significantly greater than those prepared with PEG-DMG (+PEG-DMG) (p < 0.0001 for blank LNPs and p < 0.001 for ATP-LNPs) **(Figure 1a)**. We concluded that the inclusion of ATP did not have an additional stabilizing effect on the LNP particle diameters as both ATP-loaded and blank LNPs showed similar initial diameters and as well as seven days post-storage. While ATP loading or the lack of it did not have an effect on particle diameters, we have previously demonstrated that the inclusion of siRNA, an anionic macromolecule, resulted in a dramatic stabilizing effect (lower sizes of 120 nm) as compared to blank LNPs (250 nm) (p<0.0001) (39). The observed differences in siRNA- vs. ATP-loaded LNPs can be attributed to the polyionic nature of siRNA vs. ATP and also due to the larger molecular mass of siRNA in comparison to the small molecule, ATP (13,300 *vs*. 507.18 g/mol).

**Figure 1.**
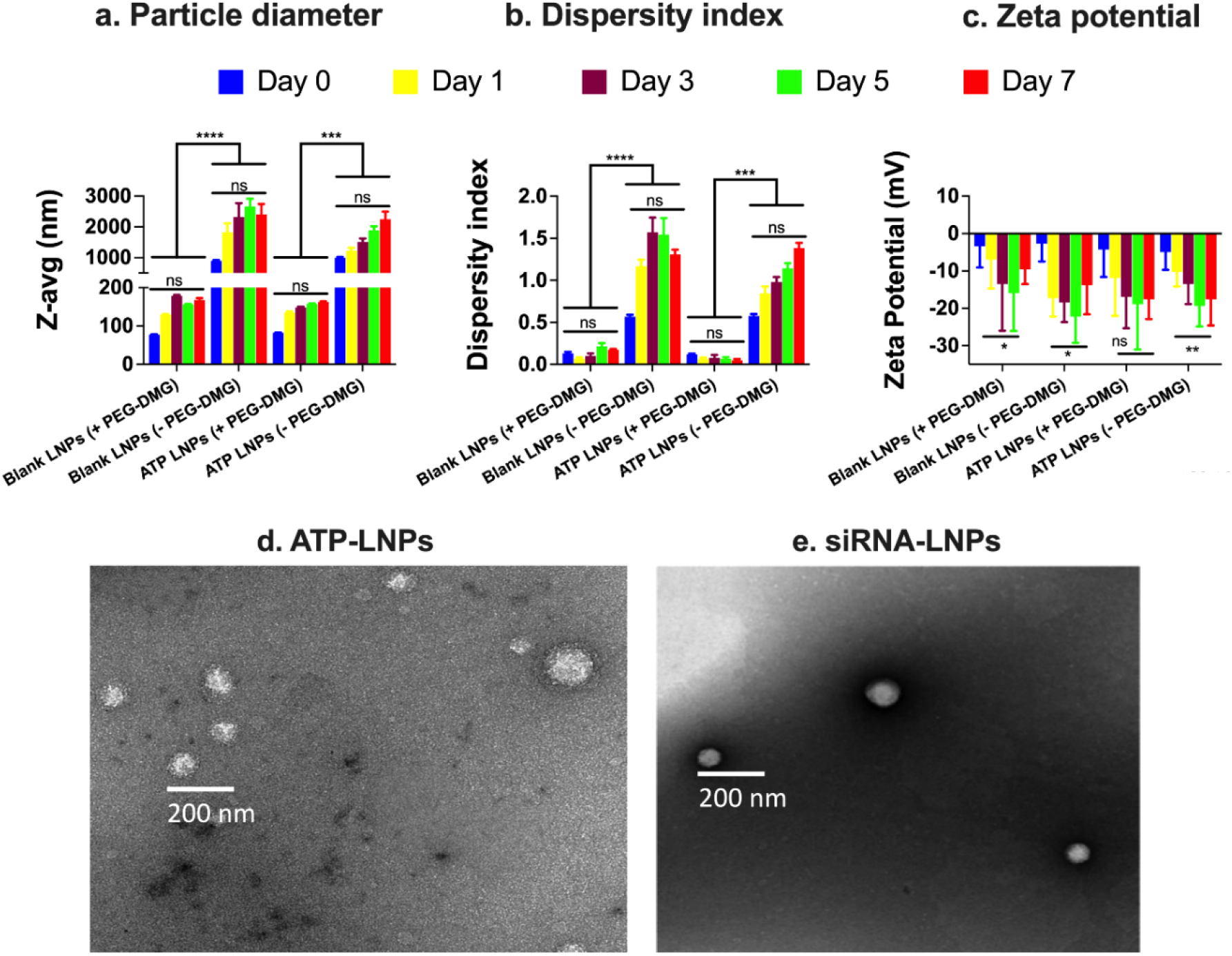
Physicochemical characterization of LNPs using dynamic light scattering and transmission electron microscopy. Blank or ATP-loaded LNPs were prepared as described in **Table 1** and siRNA-LNPs were prepared as described in Supplementary table 1. ATP-LNPs were prepared and diluted to a final ATP concentration of 5 mM using 1*x* PBS (pH 7.4). siRNA-LNPs were initially prepared in 10 mM citrate buffer and diluted to a final siRNA concentration of 50 nM using 1*x* PBS (pH 7.4). Particle diameters (**a** and **b**) and zeta potentials (**c**) were measured in 1*x* PBS and de-ionized water, respectively, using a Malvern Zetasizer Pro. Data are represented as mean ± SD of at least n=3 measurements. Negative-stain electron micrographs of ATP-LNPs (**d**) and siRNA-LNPs (**e**) were acquired using a Thermofisher TF20 electron microscope. Tukey’s and Šídák’s multiple comparisons tests were used for comparative analysis using two-way ANOVA. *p<0.05, **p<0.01, ***p < 0.001, ****p < 0.0001 and ns: non-significant.

We found a similar trend for the dispersity indices where the LNPs with PEG-DMG resulted in lower dispersity indices as compared to LNPs without PEG-DMG. The dispersity index for blank LNPs (+PEG-DMG) was found to be 0.13 that increased to 0.17 whereas the dispersity index for blank LNPs (-PEG-DMG) was 0.56 that increased to a high value of 1.3 seven days post-storage **(Figure 1b)**. The dispersity index of ATP-LNPs (+PEG-DMG) was 0.06 that decreased to 0.04 whereas that of ATP-LNPs (-PEG-DMG) was 0.57 that increased to 1.38 seven days post-storage **(Figure 1b)**. We did not find a significant difference in the dispersity indices of both blank and ATP-LNPs over a period of seven days. Dispersity values of <0.1 for ATP-LNPs (+PEG-DMG) indicate a highly monodisperse sample whereas values >1 for blank and ATP-LNPs (-PEG-DMG) indicate a polydisperse sample. There was a significant difference in the dispersity indices of blank LNPs (+PEG-DMG) *vs*. blank-LNPs (-PEG-DMG) (p<0.0001) and for ATP-LNPs (+PEG-DMG) *vs*. ATP-LNPs (-PEG-DMG) (p<0.001) **(Figure 1b)**. Thus, the inclusion of PEG-DMG was crucial in maintaining the monodisperse behavior of both blank and ATP-LNPs.

We also studied the surface charge on the LNPs by measuring their zeta potential at a pH of 7.4. The zeta potential values for all the samples ranged between -5 to -25 mV **(Figure 1c)**. Despite the negative zeta potential values, it should be noted that the samples are not strongly/solely stabilized by electrostatic interactions since all the values were < -30 mV (47). We next studied the morphology of ATP- and siRNA-LNPs under a transmission electron microscope. siRNA-LNPs were imaged at a final siRNA concentration of 50 nM whereas ATP-LNPs were diluted to 5 mM ATP. **Figures 1d and e** are representative TEM micrographs showing sphere-like morphologies of ATP- and siRNA-LNPs.

We next studied the colloidal stability of blank and ATP-loaded LNPs prepared +/- PEG-DMG upon storage at 37 °C for 24 h **(Figure 2)**. Temperature has an appreciable effect on the colloidal stability of nanoparticles and higher temperatures can potentially trigger aggregation and instability resulting in reduced *in vitro* and *in vivo* efficacy (44). While the colloidal stability of siRNA-based LNP systems is extensively studied (44), stability of LNPs encapsulating a small molecule like ATP is unknown and so, we compared the physicochemical characteristics of freshly prepared LNPs and LNPs stored at 37 °C for 24 h. The average particle diameters of blank and ATP-LNPs (+PEG-DMG) were 166 nm and 99 nm at 0 h, respectively, and 128 nm and 113 nm when measured post 24 h with interim storage at 37 °C **(Figure 2a)**. The average particle diameters of blank and ATP-LNPs (-PEG-DMG) were 496 nm and 953 nm at 0 h, respectively and increased to 1668 nm and 2811 nm, respectively, when measured post-24 h **(Figure 2a)**. We found a non-significant change/increase in the particle diameters of blank and ATP-LNPs prepared -/+ PEG-DMG when measured 24 h post-preparation. Although we did not find a significant change in LNP particle diameters, we found a significant difference (p = 0.003) in the dispersity indices of LNPs prepared without the incorporation of PEG-DMG compared to LNPs prepared with the inclusion of PEG-DMG.

**Figure 2.**
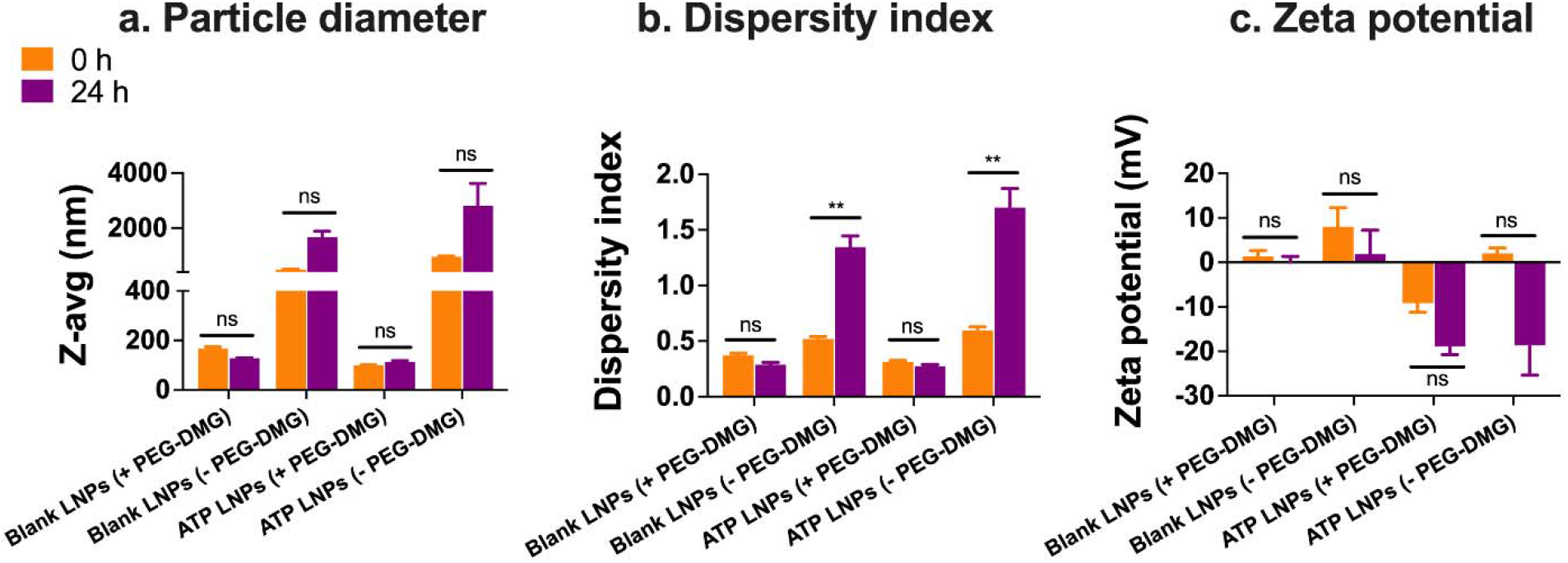
Colloidal stability of blank- or ATP-loaded LNPs immediately post-preparation (0 h) and after storage for 24 h at 37°C (24 h) studied using dynamic light scattering. ATP-LNPs were diluted to a final ATP concentration of 5 mM using 1*x* PBS (pH 7.4). Particle diameters (**a** and **b**) and zeta potentials (**c**) were measured in 1*x* PBS and de-ionized water, respectively, using a Malvern Zetasizer Pro. Data are represented as mean ± SD of at least n=3 measurements. Tukey’s and Šídák’s multiple comparisons tests were used for comparative analysis using two-way ANOVA. **p < 0.01 and ns: non-significant.

Dispersity index is a measure of particle size heterogeneity and therefore the presence of large aggregates can critically affect LNP colloidal stability. Dispersity indices below 0.3 are generally acceptable for lipid-based nanocarriers like LNPs representing a narrow, monomodal sample (48, 49). Dispersity indices of blank and ATP-LNPs (+PEG-DMG) were found to be 0.37 and 0.31, respectively, when measured immediately post-preparation and values decreased to 0.28 and 0.27 post-24 h storage **(Figure 2b)**. The average dispersity index for blank and ATP-LNPs (-PEG-DMG) were 0.51 and 0.59, respectively, that increased significantly post storage for 24 h to 1.34 and 1.7, respectively **(Figure 2b)**. These observations demonstrate a notable effect of PEG-DMG in maintaining the colloidal stability of blank and ATP-loaded LNPs. We did not notice a significant effect of ATP inclusion in the LNPs in imparting additional stability. The zeta potentials of all the samples ranged between +10 to -20 mV **(Figure 2c)**. Also, we did not note any significant difference in the zeta potentials throughout the testing regime **(Figure 2c)**.

### Serum stability of LNPs

In this experiment, we measured LNP particle diameters in the presence of 10% fetal bovine serum (FBS) to determine the stability of LNPs in cell culture conditions. We incubated blank and ATP-loaded LNPs (+/- PEG-DMG) in 10% FBS for 24 h at 37 °C. We then analyzed intensity size distribution plots to determine the effect of FBS on particle diameters. Serum proteins are known to attach on the LNP surfaces and so, we anticipated a minor difference in the sizes of LNPs containing 10% FBS (50, 51). We compared the intensity size distribution plots of LNPs either in PBS **(Figure 3, pink)** or in 10% FBS **(Figure 3, green)** immediately post-preparation **(Figure 3a)** and upon storage for 24 h at 37°C **(Figure 3b)**. We also acquired the intensity size distribution plots of a control sample containing 10% FBS in PBS to determine the size distribution and the Z-average particle diameter (Z_avg_) of serum proteins **(Figure 3, blue)**. The Z_avg_ for 10% FBS was 21 nm and 24 nm at the 0 and 24 h time points, respectively. We observed a bimodal peak for the serum proteins similar to previous reports (52-55). We looked for the presence of distinct peaks in the intensity distribution plots corresponding to those of 10% FBS (bimodal) and the LNPs. If the serum proteins destabilize and result in LNP aggregation, we expected the LNP peak to shift to larger diameters. If the LNPs are not able to maintain their intact structure in the presence of serum proteins, it is likely that the relative peak intensity of LNPs would change/decrease.

**Figure 3.**
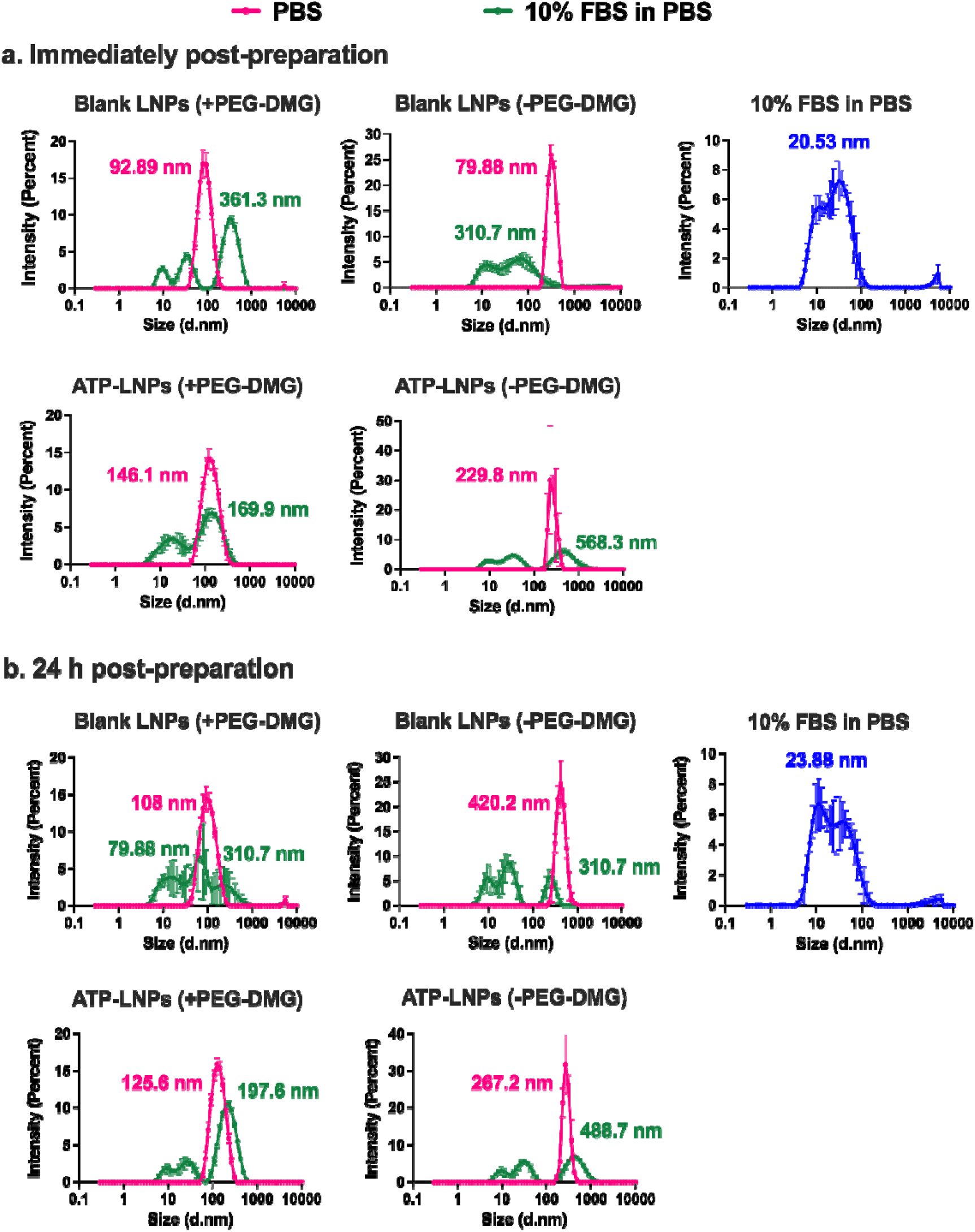
Intensity size distribution plots demonstrating the effects of 10% FBS on the stability of blank and ATP-loaded LNPs (+/-PEG-DMG) upon storage at 37 °C for 24 h. Blank and ATP-loaded LNPs +/- PEG-DMG were diluted to a final ATP concentration of 5 mM in either 1*x* PBS (pink) or 10% FBS (green). A blank sample consisting of 10% FBS in 1*x* PBS was used to profile the size distribution of serum (blue). Intensity size distributions of LNPs were obtained immediately post-preparation (**a**) and upon storage at 37 °C for 24 h (**b**). All measurements were done on a Malvern Zetasizer Pro and the data are represented as mean ± SD of at least n=3 measurements.

We overlaid the intensity distribution graphs of LNPs in PBS **(Figure 3, pink)** and LNPs in 10% FBS **(Figure 3, pink)** for the 0 h **(Figure 3a)** and 24 h **(Figure 3b)** time points. Blank LNPs (+PEG-DMG) in PBS showed a distinct peak at 93 nm that shifted to 361 nm when supplemented with FBS at the 0 h time point **(Figure 3a)**. Post-24 h, we noted peaks at 80 nm and 311 nm although at a lower intensity **(Figure 3b)** compared to the original diameter of 180 nm (in PBS). For blank LNPs (-PEG-DMG), we observed a sharp peak at 80 nm that overlapped with the bimodal peak for serum proteins when the LNPs were in 10% FBS resulting in a final Z_avg_ of 311 nm **(Figure 3a)**. The lack of a distinct peak corresponding to LNPs suggested that serum proteins may have destabilized them due to the absence of PEG-DMG in the LNP formulation. However, post-24 h we still noted the peak at 310.7 nm although with a lower %intensity compared to the peak at 420 nm for LNPs in PBS **(Figure 3b)**.

Next, we examined whether ATP loading in the LNPs may affect particle stability in the presence of serum proteins. ATP-LNPs (+PEG-DMG) in PBS showed a distinct peak at 146 nm that shifted to 169 nm when the particles were supplemented with FBS at 0 h **(Figure 3a)**. Post-24 h, we observed a sharp peak of LNPs in FBS at 198 nm that was comparable to that of LNPs in PBS with a Z_avg_ of 126 nm **(Figure 3b)**. The shift in the diameter of the blank LNP (+PEG-DMG) counterpart was more drastic along with a shift in the LNP peak as noted post-24 h in the presence of FBS when compared to ATP-LNPs (+PEG-DMG) **(Figure 3a and b)**. This suggests that ATP loading plays a notable role in maintaining the colloidal stability of LNPs in the presence of serum proteins.

ATP-LNPs (-PEG-DMG) in PBS showed a distinct peak at 230 nm, however, the peak intensity decreased along with the shift to a higher Z_avg_ of 568 nm in 10% FBS when measured immediately post-preparation **(Figure 3a)**. Twenty four hours-post storage at 37°C, LNPs in PBS showed a peak at 267 nm and LNPs supplemented with FBS retained the peak albeit at a lower intensity and a Z_avg_ of 489 nm **(Figure 3b)**. To summarize, although blank LNPs (+PEG-DMG) show a peak in the presence of FBS at 0 h, the peak was not retained at the end of 24 h incubation **(Figure 3a and b)**. In the case of blank LNPs (-PEG-DMG), we did not observe a peak at 0 h but post-24 h we noted a peak with a lower intensity. With respect to ATP-LNPs (+PEG-DMG) in FBS, we observed a sharp peak at the 0 h and 24 h time points with comparable Z_avg_ values indicating the combined stabilizing effects of ATP loading and PEG-DMG in the formulation **(Figures 3a and b)**. With ATP-LNPs (-PEG-DMG) we observed a distinct peak albeit at a lower intensity immediately post-preparation as well as 24 h after incubation **(Figures 3a and b)**. Overall, our results demonstrate that both ATP loading and PEG-DMG are crucial for maintaining the serum stability of LNPs. Interestingly, ATP loading has a more pronounced effect in maintaining LNP colloidal stability in the presence of serum proteins when compared to the inclusion of PEG-DMG.

### Flow cytometry analysis of ATP uptake into hCMEC/D3 BECs

During stroke, BECs suffer oxygen-glucose deprivation (OGD) due to decreased blood supply that affects their mitochondrial function leading to ATP depletion. We wanted to determine whether the ATP-loaded LNPs can be internalized into BECs. We used flow cytometry to quantify the cellular uptake of LNPs loaded with Alexa Fluor 647 (AF647)-tagged ATP. We spiked unlabeled ATP with AF647-ATP at a ratio of 100:1 prior to LNP preparation. For this reason, we express cellular uptake as a -fold increase in uptake over untreated cells instead of an absolute percentage (%) uptake—it should be noted that AF647-ATP was used to spike the LNPs and the LNPs also contain unlabeled ATP. We incubated the cells with samples containing either ATP-LNPs prepared w+/- PEG-DMG or free ATP or Lipofectamine-ATP complexes. We used lipofectamine as a positive control as this cationic lipid is widely used for the delivery of anionic polynucleotides such as siRNA, plasmid DNA, etc. We compared the uptake of AF647-LNPs into normoxic BECs as well as into BECs exposed to OGD as an *in vitro* model of cerebral ischemia. BECs were pre-exposed to OGD medium in a Billups-Rothenberg chamber to induce ischemic damage while the normoxic cells were maintained in complete growth medium in a normoxic incubator.

We performed a live/dead assay to account for any cell debris or dead cells and noted that about 98-99% cells were viable **(Supplementary figures 1 and 2)**. We next gated to eliminate the autofluorescence of untreated/unstained cells **(Supplementary figures 1 and 2, Black dot plot)**. A shift of the cell population to the right of the gated population indicated AF647-positive cells (AF647 (+)). Normoxic BECs incubated with free ATP showed a 2.9-fold increase in ATP uptake **(Figure 4a)** suggesting that free ATP diffuses into cells. Larger, polydisperse (995 nm, DI 0.5) ATP-LNPs (-PEG-DMG) resulted in a 3.1-fold increase in the ATP uptake by cells **(Figure 4a)**, demonstrating uptake levels similar to that of free ATP. ATP-LNPs (+PEG-DMG) resulted in a 7.7-fold increase in the ATP uptake when compared to untreated cells **(Figure 4a)**. This increased uptake is likely explained by their near-monodisperse, smaller particle diameters (∼100 nm, DI ∼0.11). We noted a 3.6-fold increase in the ATP uptake in the case of Lipofectamine/ATP complexes **(Figure 4a)**. Interestingly, ATP-LNPs (+PEG-DMG) resulted in a significantly greater uptake (p<0.0001) compared to all treatment groups **(Figure 4a)**.

**Figure 4.**
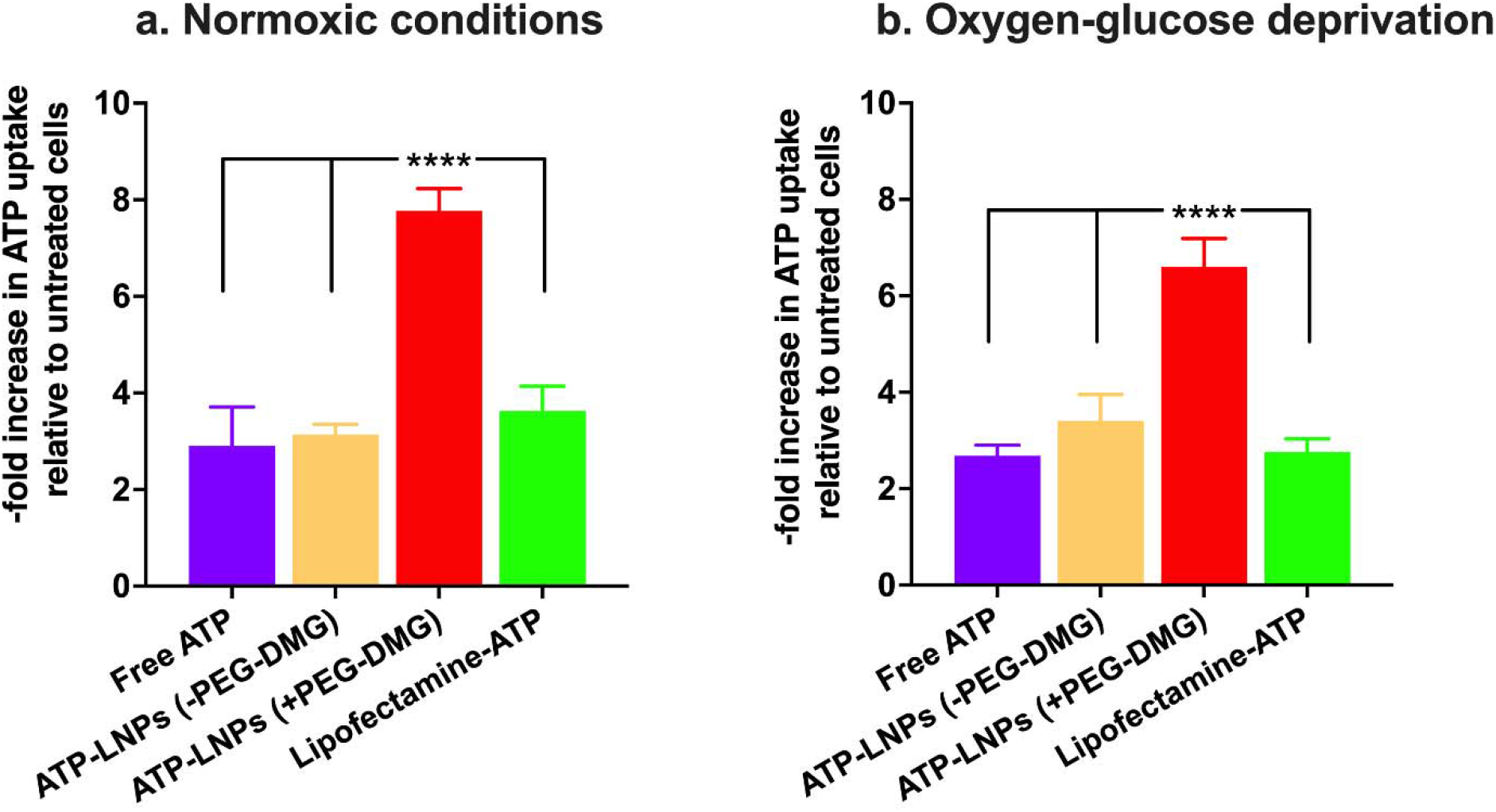
Cellular uptake of ATP-LNPs into hCMEC/D3 cells using flow cytometry. hCMEC/D3 cells were treated for 24 h with the indicated samples under normoxic conditions in a humidified incubator at 37 °C and 5% CO_2_ (**a**) under oxygen-glucose deprived conditions in the Billups-Rothenberg chamber pre-flushed with 5% carbon dioxide, 5% hydrogen and 90% nitrogen at 37 ± 0.5 °C (**b**). The treatment mixture consisted of 15 mM ATP spiked 1:100 with AF647-ATP. Lipofectamine-ATP complexes and untreated cells were used as positive and negative controls, respectively. Data are presented as -fold increase in ATP uptake (%AF647+ cells) relative to untreated cells (n=4). The raw dot plots and the gating strategy are shown in supplementary figures 3 and 4. Statistical comparisons were made with Tukey’s multiple comparisons test using one-way ANOVA. ****p < 0.0001.

We incubated hypoxic BECs with the same treatment groups. Interestingly, we observed an uptake trend that was similar to BECs cultured in normoxic conditions. We observed a 2.7-fold increase in the case of free ATP **(Figure 4b)** that was comparable to the 2.9-fold increase observed for normoxic cells **(Figure 4a)**. ATP-LNPs (-PEG-DMG) resulted in a 3.4-fold increase **(Figure 4b)** similar to the uptake noted in normoxic BECs. ATP-LNPs (+PEG-DMG) showed a 6.6-fold increase **(Figure 4b)** that was again comparable (difference is albeit non-significant with a p=0.1) to the 7.7-fold increase observed in the case of normoxic BECs **(Figure 4a)**. We observed a slightly significantly lower 2.8-fold increase in the case of Lipofectamine/ATP complexes **(Figure 4b)** (p=0.026) compared to the 3.6-fold increase in normoxic BECs **(Figure 4a)**. Overall, our results show that ATP-LNPs (+PEG-DMG) result in a significantly greater uptake of ATP into the BECs under both normoxic and hypoxic conditions (**Figure 4a and b**).

### Uptake of AF647-ATP uptake into primary human brain endothelial cells using fluorescence microscopy

Primary human brain endothelial cells (HBMECs) were incubated with samples containing 5 mM ATP spiked with AF647-ATP at a ratio of 1000:1. We speculated that LNP-delivered ATP will primarily localize in the cell cytosol as ionizable cationic lipids facilitate efficient endosomal escape (56). We incubated HBMECs with the indicated samples for 24 h followed by imaging the cells under an epifluorescent microscope (**Figure 5a**). We labeled cell nuclei using Hoechst to distinguish cytosolic AF647-ATP signals from the nuclei. **Figure 5a** shows HBMEC images under Hoechst and AF647 channels pseudo-colored as blue and green, respectively. The overlay depicts the localization of the AF647-ATP (green) fluorescence with respect to the nuclei (blue).

**Figure 5.**
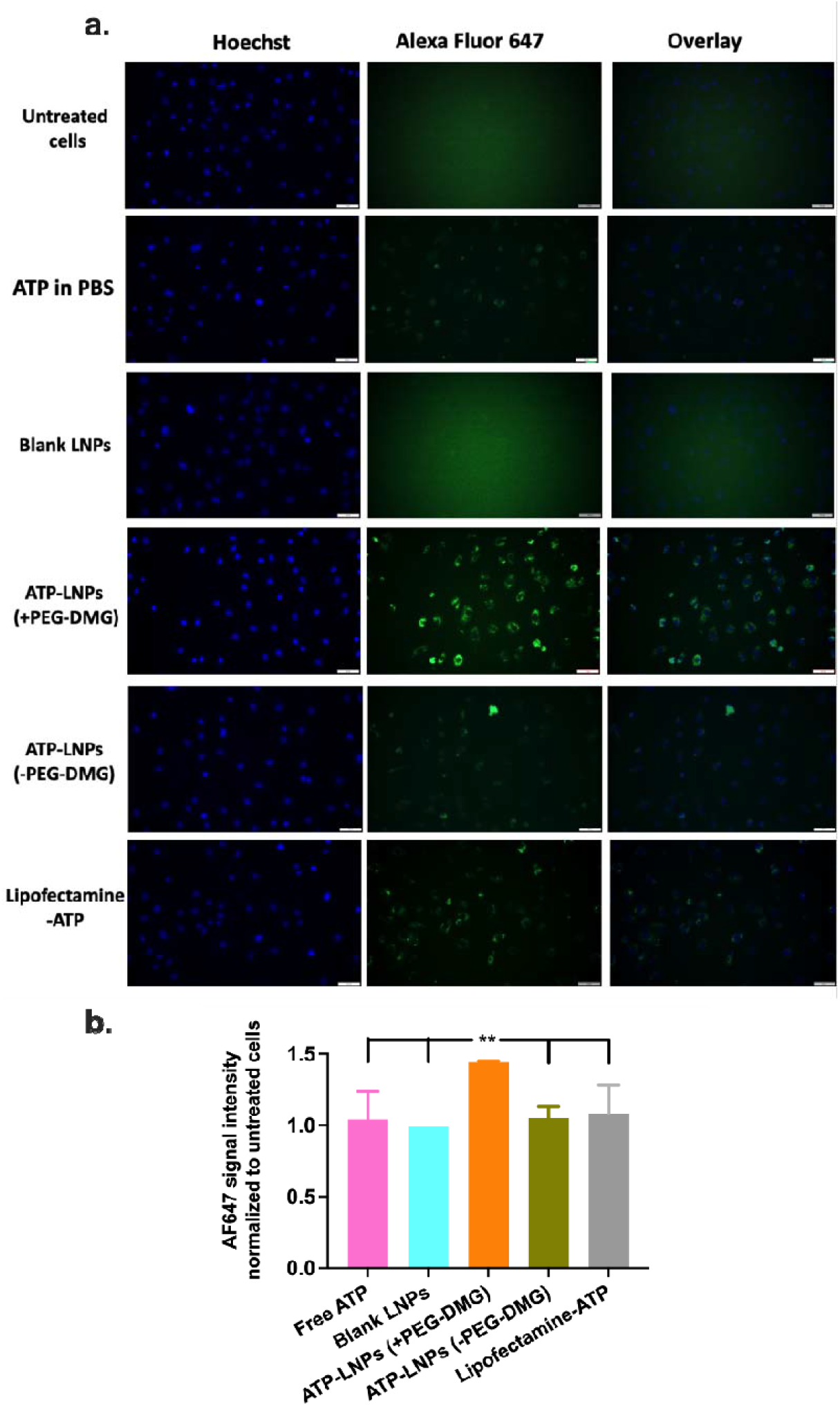
Cellular uptake of AF647-ATP into HBMECs using fluorescence microscopy. HBMECs were incubated for 24 hours with the indicated samples containing 5 mM ATP spiked 1:1000 with AF647-ATP. HBMECs were stained with 10 µg/ml Hoechst for staining the nuclei in viable cells. Scale bar = 50 µm. Cells were imaged using an Olympus IX 73 epifluorescent microscope to detect AF647 signals at excitation and emission wavelengths of 651 nm and 670 nm whereas Hoechst signals were detected at an excitation and emission wavelength of 350 nm and 461 nm, respectively. AF647 and Hoechst signals were false-colored as green and blue, respectively. Images are representative of n=4 independent wells (**a**). AF647 signals were quantified and normalized to intensities from untreated cells using CellSens Software (**b**). **p<0.01.

As seen in **Figure 5a**, we observed strong cytosolic AF647 signals (overlay images) in cells treated with AF647 ATP-LNPs (+PEG-DMG). However, cells treated with AF647 ATP-LNPs (-PEG-DMG) did not show intense AF647 signals confirming their relatively lower uptake into the cells **(Figure 5a)**. Extracellular ATP has a very short half-life of ∼10 minutes *in vivo*. Even if free ATP enters the cell by diffusion, it likely may not maintain a viable ATP concentration to achieve intracellular function. Therefore, an ideal delivery system should stably retain its cargo, i.e., ATP, extracellularly and efficiently deliver ATP intracellularly. We observed AF647 signals when the cells were treated with free/naked ATP **(Figure 5a)**. This implies that free ATP is able to diffuse in the cells however the fluorescence signals were not as intense compared to the signals noted in cells treated with LNPs (+PEG-DMG). We noted intense cytosolic AF647 signals in cells treated with Lipofectamine-ATP **(Figure 5a)**. Although Lipofectamine-ATP complexes were able to deliver ATP intracellularly, Lipofectamine has limited *in vivo* usage due to its strong cationic nature leading to toxicity (57). We also quantified fluorescence intensities of the images using CellSens software and noted a significantly higher (p<0.01) AF647 signal intensity in cells treated with ATP-LNPs (+PEG-DMG) as compared to the other treatment groups **(Figure 5b)**.

### Effects of LNP exposure on relative ATP levels in HBMECs

We studied changes in the ATP levels in HBMECs **(Supplementary figure 3)** treated under normoxic and hypoxic conditions. We incubated cells with treatment mixtures containing different concentrations of ATP for 4 h or 24 h under normoxic and hypoxic conditions **(Supplementary figures 3a and 3b)**. We treated the cells with 0, 2.5, 5, 10, 15 and 20 mM free ATP or ATP-LNPs for 4 or 24 h in complete growth medium (normoxic) or OGD medium (hypoxic). We observed an increasing trend in the relative cellular ATP levels when they were treated with the different ATP concentrations in normoxic as well as hypoxic conditions **(Supplementary figure 3)**. However, we did not note a difference in the ATP levels 24 h post-incubation compared to the 4 h incubation time. Similarly, we did not observe a difference in the ATP levels of cells when treated with free ATP or ATP-LNPs under normoxic as well as hypoxic conditions. However, we observed significantly higher ATP levels in cells under hypoxic conditions **(Supplementary figure 3b)** as compared to normoxic conditions **(Supplementary figure 3a)** for the 10, 15 and 20 mM ATP treatment concentrations (p<0.05, p<0.01 and p<0.001). This likely suggests that hypoxic BECs have a higher probability of showing increased relative ATP levels compared to normoxic cells. However, there are a few caveats associated with the use of luciferin-luciferase based assays that rely on the ability of luciferase to oxidize luciferin to oxyluciferin in the presence of ATP (58). Most importantly, we used ATP concentrations in the mM range which exceeds the 1 µM upper limit of detection of luciferin-luciferase assays (59). Therefore, we were unable to reliably quantify the effects of LNP-delivered ATP. Also, some luciferases have the ability to generate new ATP molecules from the intracellular pool of ADP already present. This results in the lack of specificity towards the ATP that is present in the sample (60). Additionally, AMP and ADP can act as competitive inhibitors of ATP, further limiting the assay specificity (61).

## Discussion

LNPs have emerged as successful non-viral carriers for the delivery of high molecular mass polynucleotides such as mRNA, siRNA and pDNA (6, 7, 62, 63). In this pilot study, we investigated the ability of LNPs to formulate a small molecule drug, ATP—a novel and previously unexplored application of LNP delivery. LNPs have shown beneficial pharmacokinetics, efficient protection of the payload and uptake into cells and lower immune activation. These characteristics make them compelling carriers for drug delivery (5, 7, 29, 31, 36-38, 40, 63, 64). Their high encapsulation efficiencies with mRNA and siRNA cargo led us to explore LNPs as a carrier for formulating ATP. We loaded ATP into the LNPs at an ionizable cationic lipid/ATP weight:weight ratio of ca. 1:1 (**Table 1**) and using a luminescence-based ATP assay, we noted similar luminescence values for free ATP vs. LNP-loaded ATP, suggesting near-complete loading of ATP into the LNPs (**Supplementary table 2**).

Cerebral ischemia/reperfusion injury results in mitochondrial dysfunction and decreases ATP production in BECs lining the BBB (65). Free ATP has a very short half-life (∼10 minutes). Here, we have formulated ATP-LNPs for ATP uptake by the BECs, whose protection is of central importance in conditions like stroke (14, 66, 67). Our prior works have shown that transfer of mitochondria to hypoxic BECs increased their relative ATP levels and concomitantly, their cell viability (14, 67). Because of their critical reliance on ATP for their metabolic function and structural integrity, we chose to deliver ATP to offset the energy imbalance in ischemic BECs.

In our previous work, we reported the effect of incorporation of PEG-DMG on the colloidal stability of siRNA-LNPs (39). We also demonstrated that the incorporation of siRNA provides an additional stabilizing effect to LNPs in addition to that conferred by PEG-DMG (39). Our previous observations indicated that both siRNA (cargo) and the PEG-DMG (stealth component) improve LNP stability on storage at 2-8 °C for a period of seven days and when supplemented with 10% FBS upon storage at 37 °C for a period of 24 h. We wanted to understand if similar factors play a role in maintaining the colloidal stability of ATP-LNPs.

First, we characterized particle diameters and surface charge of blank and ATP-loaded LNPs prepared +/- PEG-DMG over a period of seven-days upon storage at 2-8 °C. We observed a non-significant increase in the particle diameters and dispersity indices of blank and ATP-loaded LNPs prepared +/- PEG-DMG over a period of seven days **(Figures 1a and b)**. We observed a significant difference in the particle diameters and dispersity indices of blank LNPs (+PEG-DMG) *vs*. blank LNPs (-PEG-DMG) (p<0.0001) and ATP-LNPs (+PEG-DMG) *vs*. ATP-LNPs (-PEG-DMG) (p<0.001) **(Figures 1a and b)**. These findings corroborate our earlier observation that PEG-DMG is a crucial component to maintain lower particle diameters and dispersity indices of both blank and ATP-LNPs. However, we did not find a significant difference in the colloidal properties of blank *vs*. ATP-loaded LNPs **(Figures 1a and b)**. This implies that while siRNA loading played an important role in stabilizing siRNA-LNPs (39), the small molecule ATP does not have a similar effect in the case of ATP-LNPs. This can be attributed to the polyionic nature and large molecular mass of siRNA as compared to ATP. The polyionic siRNA likely forms multiple cooperative interactions with the cationic lipids while ATP, a small molecule cargo, may not form the additional cooperative bonds that likely resulted in the superior stability of siRNA-loaded LNPs.

Next, we studied the surface charge (zeta potential) of the prepared LNPs over a period of seven-days upon storage at 2-8 °C. We observed negative zeta potential values for all the samples (between -5 to -25 mV) indicative of the effects of PEG-DMG-mediated steric stabilization in combination with the loading of ATP into the LNPs. We did not observe a significant difference in the zeta potential of LNPs over a period of seven-days for ATP-LNPs (+PEG-DMG) suggestive of their stability. However, we did observe a significant difference in the zeta potentials of blank LNPs (+/- PEG-DMG) (p<0.05) and ATP-LNPs (-PEG-DMG) (p<0.01) **(Figure 1c)**. We next studied the morphology of ATP-LNPs using TEM and compared it to the morphology of siRNA-LNPs that has been extensively reported (64, 68). siRNA-LNPs show a spherical morphology with a ring structure enclosing an amorphous core as previously reported (64). We noted a similar morphology for ATP-LNPs **(Figures 1d and e)**.

We characterized the colloidal stability of LNPs post-storage for 24 h at 37 °C. We did not observe a significant increase in the particle diameters and zeta potential values of the ATP-LNPs (+PEG-DMG) over a period of 24 h **(Figures 2a and c)**. Also, we did not note a significant difference in the particle sizes of blank LNPs (+PEG-DMG) *vs*. blank LNPs (-PEG-DMG) or ATP-LNPs (+PEG-DMG) *vs*. ATP-LNPs (-PEG-DMG) **(Figure 2a)**. For dispersity indices, we did not observe significant differences among any given sample, however, we did note a significant difference for blank LNPs (-PEG-DMG) and ATP-LNPs (-PEG-DMG) when comparing 0 h (immediately after preparation) *vs*. 24 h-post preparation (p<0.01) **(Figure 2b)**. Both blank and ATP-loaded LNPs prepared in the presence of PEG-DMG showed improved stability over the 24 h period, suggestive of steric stabilization (46). Ball *et al*. determined the effect of extreme freeze-thaw cycles and pH conditions on the colloidal stability and *in vitro* gene silencing efficacy of siRNA-LNP systems over a period of 10 months (44). The authors studied changes in the Z_avg_ and dispersity indices of the LNPs over pH ranging from 3 through 9 at -20, 2 and 25 °C. Storage of siRNA-LNPs at -20°C in pH 3-9 increased particle diameters and DI values from ca. 200 nm to >500 nm and from 0.1 to >0.5, respectively. The increased particle diameters and DI values of siRNA-LNPs concurrently decreased their *in vitro* gene silencing efficacy by 8 to 10-fold (44).

Stability of the prepared LNPs was also tested in cell culture conditions using LNPs diluted in 10% FBS immediately post-preparation (0 h) and 24 h post-preparation at 37°C **(Figure 3 a and b)**. We compared the intensity distribution peaks of LNPs in 1*x* PBS *vs*. 10% FBS to determine the effects of serum on particle diameters and their resulting stability. We observed that the LNP peak was retained only in the ATP-LNPs (+PEG-DMG) sample in serum conditions **(Figure 3)**. For the other samples i.e., blank LNPs (+/-PEG-DMG) and ATP-LNPs (-PEG-DMG) the LNP peak was not retained, or the peak overlapped with the peak corresponding to the serum proteins **(Figure 3)**. These observations suggest that the inclusion of ATP played a critical role in maintaining particle diameters in the presence of serum proteins although it did not seem to have an effect in 1*x* PBS (**Figure 3**). Overall, we conclude that the inclusion of both ATP and PEG-DMG play an important role in the stabilization of LNPs in the presence of serum proteins.

Next we used flow cytometry and fluorescence microscopy to quantify the uptake of ATP spiked with AF647-ATP at ratios of 100:1 and 1000:1, respectively. Flow cytometry data revealed a significantly higher uptake of AF647-ATP by normoxic (7.7-fold) **(Figure 4a)** as well as hypoxic (6.6-fold) BECs when delivered via LNPs (+PEG-DMG) (p<0.0001) **(Figure 4b)**. This suggests that the superior colloidal stability of ATP-LNPs (+PEG-DMG) facilitates their improved uptake into BECs. We then studied whether the internalized ATP taken is localized primarily in the cell cytosol as a result of endosomal escape. Fluorescence microscopy was used to image cells treated with AF647 ATP-spiked LNPs. Similar to the flow cytometry results, we observed strong AF647 signals in the cytoplasm of cells treated with ATP-LNPs (+PEG-DMG) **(Figure 5)**. AF647 signal was observed specifically in the cell cytosol (green AF647 signals surrounding the nuclei) with the greatest intensity noted in the cells treated with ATP-LNPs (+PEG-DMG) **(Figure 5)**. Thus, normoxic as well as hypoxic cells internalized ATP when treated with ATP-LNPs (+PEG-DMG), and the uptake was superior when compared to free ATP, ATP-LNPs (-PEG-DMG) and Lipofectamine-ATP complexes **(Figure 5)**. LNPs formulated with ionizable cationic lipids allows efficient endosomal escape followed by cytosolic release of the cargo. In the acidic environment of the endosome, the ionizable cationic lipid attains a positive charge and associates with the negatively charged endosomal lipids causing disruption of the endosomal membrane and cargo release into the cell cytoplasm (30, 69). Our data suggests that LNPs (+PEG-DMG) deliver ATP into the cytosol.

The loss of mitochondrial ATP triggers the ionic, biochemical and cellular imbalances in ischemic stroke (65). Protection of BECs is critical for maintaining brain homeostasis as they maintain functional interactions with astrocytes and other support cells to form the BBB, the neurovascular unit and the neural vascular niche (70). The BECs lining the BBB are an accessible target for ischemic protection as opposed to the brain parenchyma tissue behind the BBB (66). Hypoxia affects pinocytosis in BECs, causes higher cellular volumes, induces structural changes in tight junction proteins and decreases the metabolic activity of BECs (71-74). Despite these cellular alterations, LNPs were internalized into hypoxic BECs. Overall, our results suggest the potential of LNPs to deliver small molecule actives such as ATP to protect the ischemic BBB.

## Conclusions

For the first time, we formulated ATP, a small molecule compound using the clinically-successful LNP platform. ATP-LNPs formulated in the presence of PEG-DMG showed optimum colloidal stability at 2-8 ºC, 37 ºC and in the presence of serum proteins. Their improved colloidal stability translated to their ability to increase the intracellular uptake of ATP into BECs, a low pinocytic cell model. Our results demonstrate the potential of LNPs to deliver anionic small molecules such as ATP to BECs lining the BBB.

## Acknowledgements

This work was supported via start-up funds for the Manickam laboratory from Duquesne University (DU), a 2021 Faculty Development Fund (Office of Research, DU) and the Charles Henry Leach II Fund to the PI. The authors express their deep appreciation to Mr. Kandarp Dave for experimental guidance. The authors are thankful to Dr. Lauren O’Donnell, and Mss. Manisha Chandwani and Yashika Kamte (DU) for flow cytometry support.

