## Supplemental Files for "Lipidoid nanoparticles increase ATP uptake into hypoxic brain endothelial cells"

**Formulation of siRNA-LNPs**

For siRNA-LNPs, a one mg/mL solution of siRNA was prepared in 10 mM citrate buffer, pH 4.0. The LNPs were then prepared according to the scheme shown in ***Supplementary table* *1*** using the ‘fast mixing’ method as previously described (1). siRNA-LNPs were made at a final siRNA concentration of 400 nM, unless stated otherwise.

**Supplementary table 1. Representative formulation scheme for siRNA- LNPs**

| Component | w/w% in final LNP mixture | Stock concentration (mg/mL) | Volume of the stock solution required (µL) |
| --- | --- | --- | --- |
| Ethanolic phase | | | |
| C12-200 | 50 | 1 | 12.5 |
| Cholesterol | 38.5 | 1 | 3.1 |
| PEG-DMG | 1.5 | 0.5 | 13.6 |
| DSPC | 10 | 0.5 | 16.9 |
| Ethanol | - | - | 78.9 |
| Total ethanolic phase volume | | | 125 |
| Aqueous phase | | | |
| siRNA | 400 nM | 1 | 2.5 |
| 10 mM citrate buffer | - | - | 122.5 |
| PBS | - | - | 250 |
| Total aqueous phase volume | | | 375 |
| Final volume of LNPs prepared (µL) |  |  | 500 |

**Supplementary table 2. Luminescence values for free ATP and ATP-LNPs**

| **Groups** | **Blank LNPs (+PEG-DMG)** | **Blank LNPs**  **(-PEG-DMG)** | **ATP-loaded LNPs (+PEG-DMG)** | **ATP-loaded LNPs**  **(-PEG-DMG)** | **Free ATP in PBS** | **Blank PBS** |
| --- | --- | --- | --- | --- | --- | --- |
| **Average luminescence (A.U.)** | 1681.3 | 3236.7 | 70260.3 | 70375.3 | 60735.7 | 158.0 |

Luminescence values for the indicated samples were obtained using the Cell Titer Glo ATP assay described below (*see* Effect of LNPs on resulting ATP levels in HBMECs).

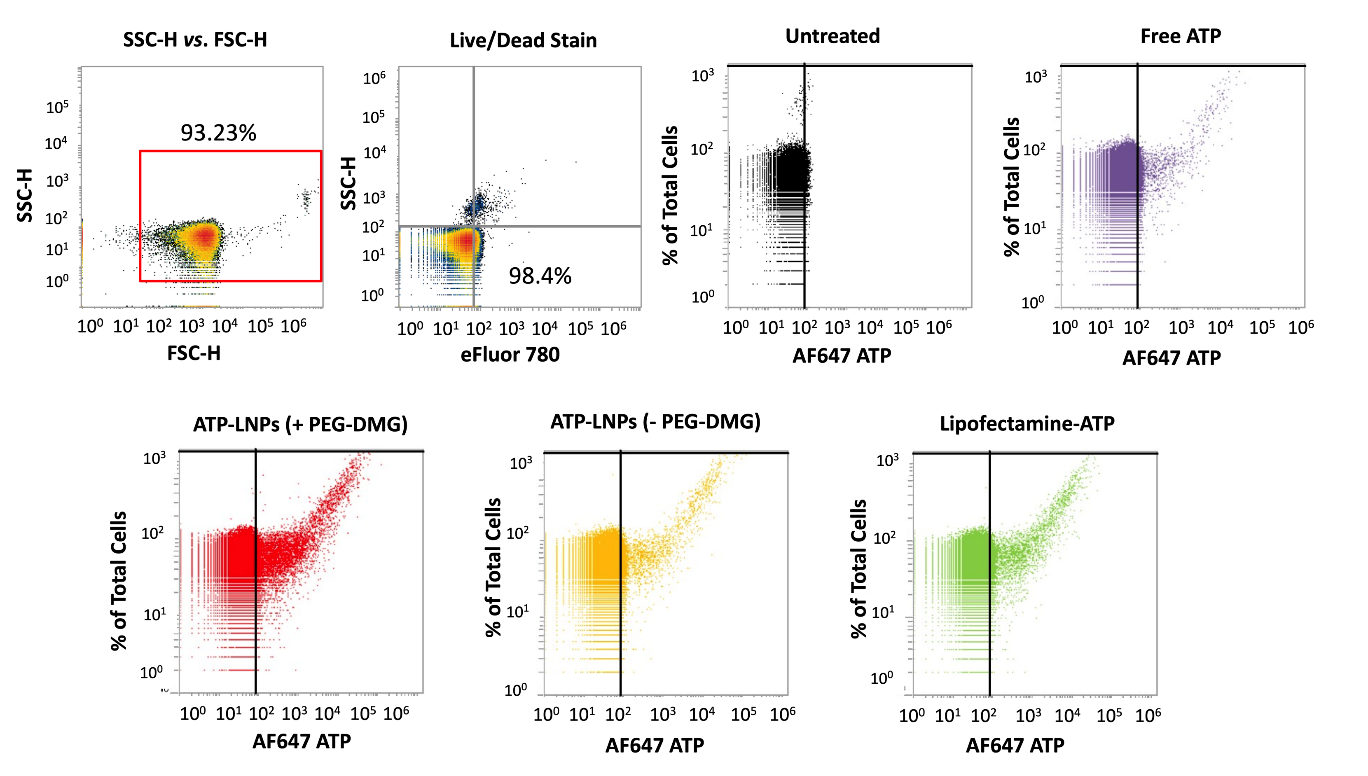

**Supplementary figure 1. Uptake of AF647-ATP into normoxic hCMEC/D3 cells using flow cytometry analysis.** Cells were maintained under normoxic conditions in complete growth medium in a humidified incubator at 37 °C and 5% CO_2_. Cells were treated for 24 h with the indicated samples containing 15 mM ATP spiked 1:100 with AF647-ATP. Lipofectamine-ATP complexes and untreated cells were used as positive and negative controls, respectively. Data are presented AF647 (+) cells noted by the shift past the solid black line after gating out the autofluorescence of the untreated cells.. The dot plots are representative of quadruplicate samples.

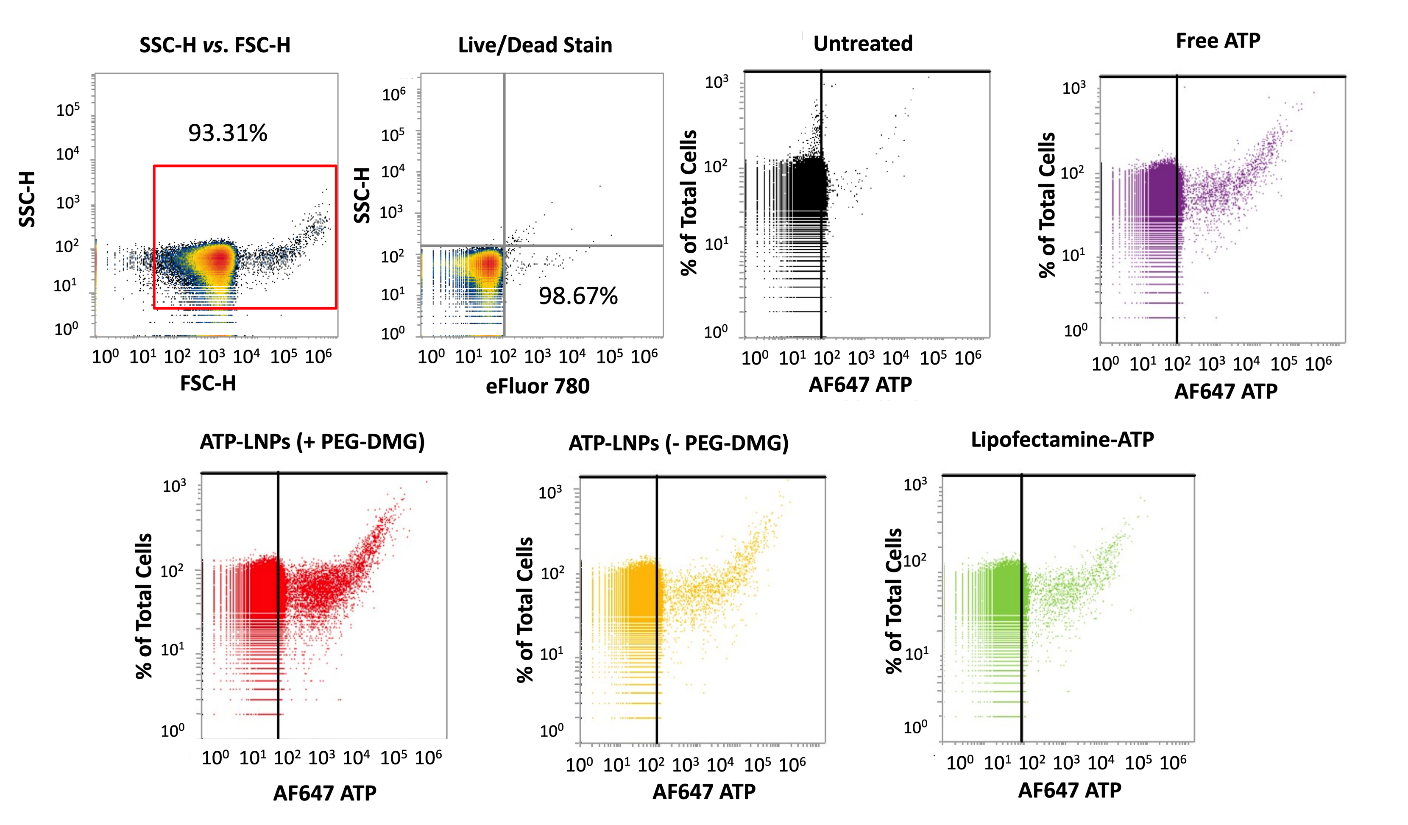

**Supplementary figure 2. Uptake of AF647-ATP into hypoxic hCMEC/D3 cells using flow cytometry analysis.** Cells were maintained under hypoxic conditions using OGD medium in the Billups-Rothenberg chamber pre-flushed with 5% carbon dioxide, 5% hydrogen and 90% nitrogen at 37 ± 0.5 °C. Cells were treated for 24 h with the indicated samples containing 15 mM ATP spiked 1:100 with AF647-ATP. Lipofectamine-ATP complexes and untreated cells were used as positive and negative controls, respectively. Data are presented AF647 (+) cells noted by the shift past the solid black line after gating out the autofluorescence of the untreated cells. The dot plots are representative of quadruplicate samples.

**Effect of LNPs on resulting ATP levels in HBMECs**

HBMECs were seeded in a cell attachment factor treated clear, flat bottom Poly-D-Lysine coated 96-well plate (Azer Scientific, Morgantown, PA) at a density of 16,500 cells/well in complete growth medium. Untreated cells and cells treated with 100 μg/mL polyethylene imine (PEI) were used as controls. Treatment groups comprised of the indicated concentrations of either free ATP or ATP-LNPs (**Supplementary figure 3**). Cells were treated with the respective groups for 4 h and 24 h in complete growth medium (normoxic) or in OGD medium (hypoxic) in a final volume of 50 μL/well. Cells were exposed to hypoxic conditions via pre-incubating them in OGD medium in the Billups-Rothenberg chamber for 24 h. Post-24 h of OGD exposure, cells were exposed to the indicated samples in either complete growth medium (normoxic) or OGD medium (hypoxic) in the normoxic incubator. The cells were then washed with 1*x* PBS to remove any extraneous ATP as a result of the treatment. Sixty μL of complete growth medium and 60 μL of the Cell Titer-Glo reagent were added to each well of a 96-well plate. The plate was incubated in dark for 15 minutes in an incubator-shaker (Thermo Fisher Scientific, Waltham, MA) at room temperature. Sixty μL of the mixture from each well was transferred to a flat bottom, white opaque 96-well plate (Azer Scientific, Morgantown, PA) and luminescence was measured using SYNERGY HTX multi-mode reader (BioTek Instruments, Winooski, VT). The relative ATP levels (%) was normalized to the untreated cells as shown in **Equation 1.**

Relative ATP levels (%) = $\frac{Luminescence from cells treated with samples}{Luminescence from untreated cells}\times100$ **Equation 1**

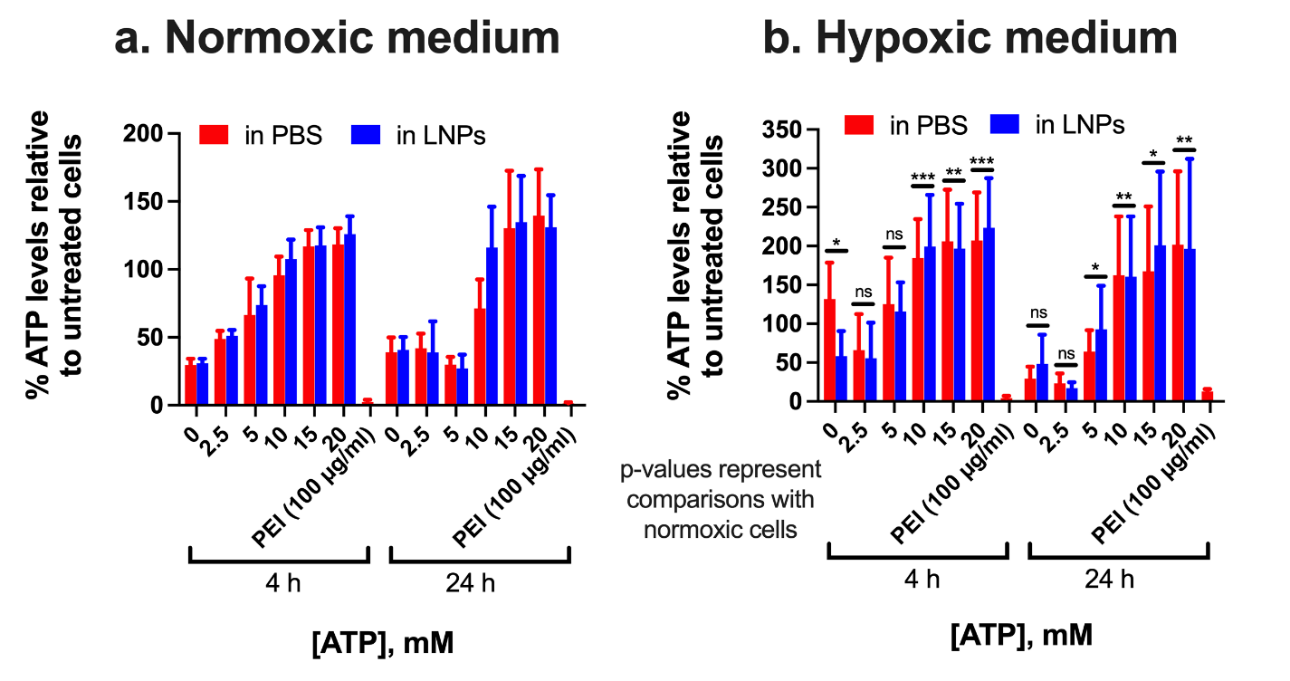

**Supplementary figure 3.** **Effect of LNPs on relative ATP levels in HBMECs under normoxic vs. hypoxic conditions.** Primary HBMECs were maintained for 24 h in either normoxic medium or in OGD medium in the Billups-Rothenberg chamber. The cells were then treated with indicated samples for 4 h or 24 h in normoxic or hypoxic medium. Untreated cells and cells treated with PEI (100 μg/mL) were used as negative and positive controls, respectively. % ATP levels were determined post-treatment using a CellGlo luminescence assay, and the data was normalized to untreated cells. Data represents mean + SD (n=6). Tukey’s and Šídák's multiple comparisons tests for statistical comparisons were performed using two-way ANOVA. **P-values represent comparisons between ATP levels of hypoxic cells vs. normoxic cells.** *p<0.05, **p < 0.01, ***p < 0.001 and ns: non-significant.
